# Quantitative computerized analysis demonstrates strongly compartmentalized tissue deformation patterns underlying mammalian heart tube formation

**DOI:** 10.1101/2025.08.03.668375

**Authors:** Morena Raiola, Miquel Sendra, Jorge Nicolás Dominguez, Miguel Torres

**Affiliations:** Cardiovascular Regeneration Program, Centro Nacional de Investigaciones Cardiovasculares (CNIC), 28029 Madrid, Spain; Department of Experimental Biology, Faculty of Experimental Sciences, University of Jaén, 23071 Jaén, Spain; Centro de Investigación Biomédica en Red de Enfermedades Cardiovasculares (CIBERCV), 28029 Madrid, Spain

**Author notes:** CNRS, UTLN, LIS 7020, Turing Centre for Living Systems, Aix Marseille University, Marseille, France.

## Abstract

The quantitative analysis of tissue deformation at cellular resolution remains an important challenge in mammalian organogenesis. Here, we developed a new computational workflow to extract regional and temporal patterns of tissue deformation and applied it to a collection of live microscopy datasets from mouse cardiogenesis. We devised a method to track tissue deformation directly from time-lapse raw images and experimentally validated the method by comparison with actual cell tracks. We then used a machine-learning approach to temporally and spatially align different specimens and reconstruct a single statistical model of tissue motion, deducing maps of strain, anisotropy, and tissue growth. We also implemented a virtual fate mapping tool that allows tracking any initial position in the cardiac primordium onto the linear heart tube. Our study reveals predominant local cellular coherence during the deformation of the cardiac tissue, whereas strong compartmentalization of tissue deformation patterns transforms the bilateral cardiac primordium into a 3D longitudinal heart tube. At the future outer curvature of the primitive tube, the ventricular chamber forms by expansion of the tissue in a hemi-barrel shape with two harnessing belts; one that constrains tissue expansion at the arterial pole and one that constrains the expansion at the venous pole. Our study provides a new approach to understanding heart morphogenesis and proposes a new model of primitive heart tube formation.

## INTRODUCTION

Quantifying tissue morphogenesis at cellular level is essential for understanding developmental mechanisms, addressing congenital heart disease, and designing tissue engineering strategies. In mammals, the heart is the first organ to form and function. Heart morphogenesis is particularly challenging to study because it involves extreme tissue deformation and occurs concurrently with the onset of cardiac beating. Early cardiogenesis involves extensive reorganization and progressive differentiation of the cardiogenic fields to rapidly create a pumping structure (Ivanovitch et al., 2017; Kelly et al., 2014; Meilhac et al., 2014; Buckingham et al., 2005; Moorman & Christoffels, 2003). In mouse, the first differentiation wave produces a simple single layer known as the cardiac crescent (CC). The CC then undergoes a series of deep deformations, gradually forming the heart tube (HT), a sophisticated three-dimensional structure with inflow (IFT) and outflow (OFT) tracts and ability to support embryonic circulation (de Boer et al., 2012). While the evolution of tissue shape during cardiogenesis has been characterized and quantitatively analyzed (Ebrahimi et al., 2022; Esteban et al., 2022; Kawahira et al., 2020; J.-F. Le Garrec et al., 2013), understanding how tissues deform during morphogenesis requires the definition of cell displacement and rearrangement patterns.

Recent advances in microscopy and genetic cell labelling have opened new avenues for investigating tissue dynamics. Live imaging techniques, enabling non-invasive observation of biological phenomena, have opened new opportunities and perspectives in this field (Dominguez et al., 2023; Raiola et al., 2023; Sendra et al., 2022; Ivanovitch et al., 2017; J. F. Le Garrec et al., 2017). These new approaches have provided access to datasets that include tissue deformation and cellular tracks; however, these datasets remain underexploited for global, multiscale analysis of tissue deformation during cardiogenesis.

While some studies have focused on the dynamic analysis of discrete morphogenetic events of early heart development (Sendra et al., 2025; Ivanovitch et al., 2017; Buckingham et al., 2005; Meilhac et al., 2004), the global dynamic analysis of tissue and cell dynamics during mammalian heart development has not been achieved. A thorough understanding of the morphogenetic mechanisms of early heart development requires adopting a multiscale approach that incorporates the behavior of individual cells, but at the same time describes the global dynamics at the tissue level.

Here, we address this challenge by applying a previously developed and validated computational workflow, described in detail in Raiola et al., 2025, to quantify and compare tissue deformation patterns by integrating motion profiles estimated from multiple 3D live fluorescence microscopy datasets. The workflow is based on a non-invasive analysis of individual live images and quantifies tissue kinematics to compute deformation fields and principal strain directions, without modelling the forces underlying such motion. To align the individual motion profiles of multiple specimens over time, the workflow incorporates a staging system that synchronizes 3D live images with a previously described pseudo-dynamic Atlas of tissue geometry during heart morphogenesis (Esteban et al., 2022). This strategy allows the deduction of deformation patterns with spatial and temporal resolution, capturing both instantaneous features and their cumulative history. In this study, we use this framework to extract and interpret biologically meaningful deformation patterns during heart development. By applying this established workflow to heart development, we report new essential features of cardiogenic tissue deformation, including a strong spatio-temporal compartmentalization of cell behaviors. The 3D+t model generated also enables *in-silico* fate map analysis at the cellular or regional level, which provides a tool to interrogate specific features of the dynamics of cardiac cells during HT morphogenesis. This model lays the groundwork for future predictive simulations of the forces driving heart morphogenesis.

## RESULTS

### Estimating Myocardial Motion and Deformation from Multiple Time-Lapse Datasets

We applied a workflow that consists of four steps summarized in Figure 1: (A) Individual time-lapse analysis to capture the geometry of the myocardium from the cardiac crescent to the linear heart tube; (B) Integration of multiple time-lapse datasets through spatiotemporal registration into a previously described 3D+t Atlas (Esteban et al., 2022); (C) Quantification of cardiac spatiotemporal deformation patterns during CC to HT morphogenesis; (D) *In-silico* fate mapping to investigate how different CC regions contribute to the forming HT. The full methodology is detailed in a co-submitted manuscript (Raiola et al., 2025).

**Figure 1:**
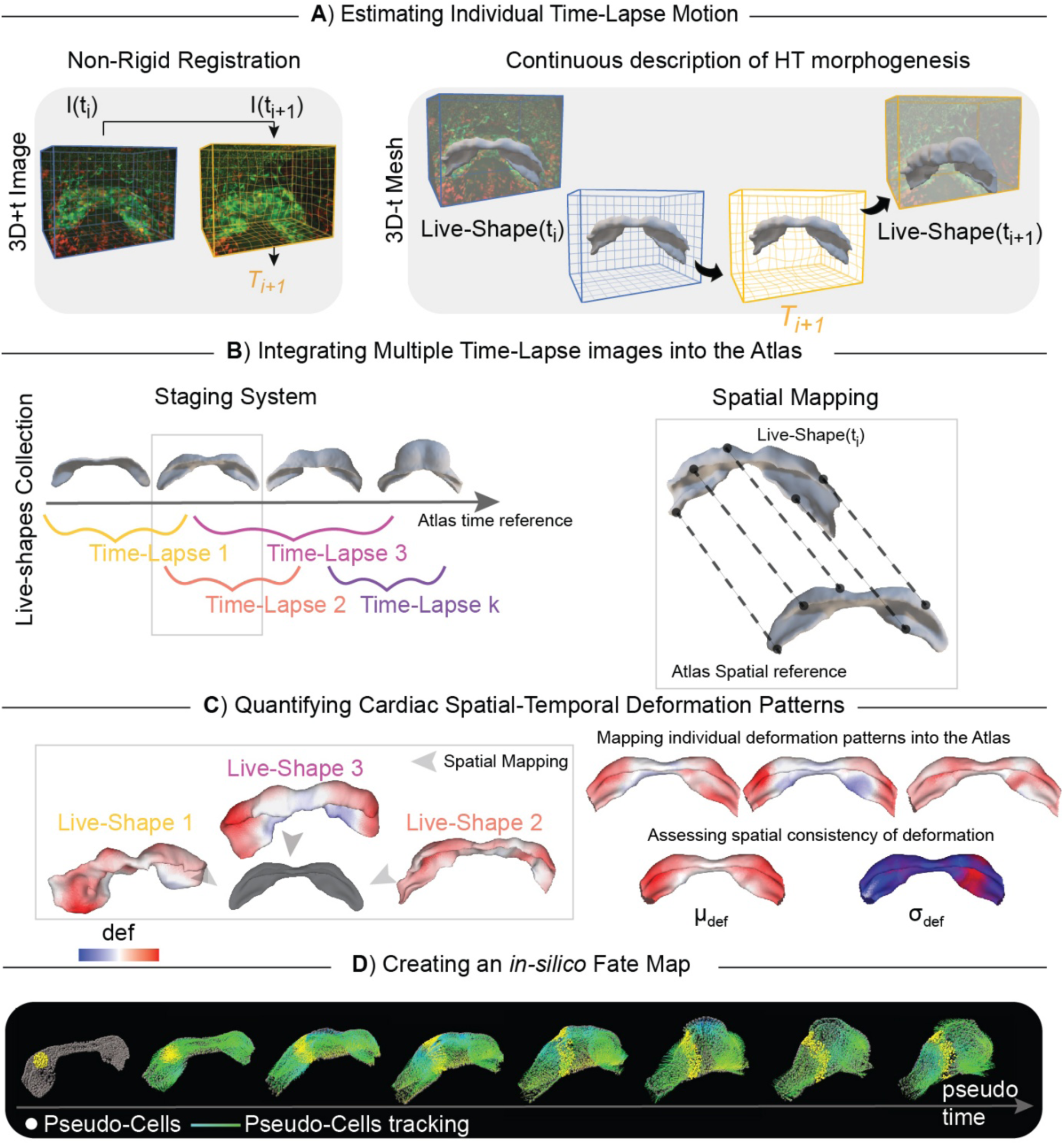
Workflow for extracting and defining the motion and deformation pattern of the myocardium during early heart morphogenesis. The analysis pipeline consists of four main steps. Our dataset includes 16 specimens ranging from E7.75 to E8.25 (12 hours). A), Estimating Individual Time-Lapse Motion: A non-rigid algorithm extracts the transformation T needed to deform one frame, I(ti), into the next frame, I(ti+1). The transformation is computed for a set of control points of a regular 3D grid (blue grid). Control point displacement is updated (orange grid) until the two frames overlap. We collect the set of transformations {Ti} for each time-lapse image to compute the continuous description of HT morphogenesis. B), Integrating multiple time-lapse images into the Atlas: We incorporate the individual continuous descriptions of HT deformation into the Atlas. We define a staging system that aligns the individual time-lapse images on a common time reference and registers the related 3D mesh (Live-Shape) onto the spatial reference (Atlas). C), Quantifying Cardiac Spatial-Temporal Deformation Patterns: We extract the deformation patterns from each continuous description of HT deformation and compile them into the Atlas, generating a single stage-by-stage model (μdef, σdef). D), Creating an in-silico Fate Map: We concatenate a continuous description of HT deformation from the individual steps to recreate a unique and continuous description of early HT deformation patterns. Gray spots represent the initial positions of the pseudo-cells, while color lines, their displacement during development. Yellow spots indicate the tracked cells.

We analyzed both published (Ivanovitch et al., 2017) and new datasets consisting of 3D+t live images (Raiola et al., 2025), including 16 embryos from early CC formation to the onset of heart looping. Myocardial cells were labelled using various reporter strategies, including Nkx2.5GFP (Wu et al., 2006) or a combination of Nkx2.5Cre (Stanley et al., 2004) with different fluorescent reporter alleles. To enable individual cell tracking, Nkx2.5GFP embryos carried sparse genetic labels induced by low-dose tamoxifen using the RERT allele (CreERT2 knocked-in under the control of RNApol2 promoter Guerra et al., 2003); these labels were used exclusively as fiducial markers for validation, not for quantitative single-cell analysis. Additional embryos expressed Mesp1Cre to mark all mesodermal cells, or Islet1Cre to label cardiac progenitor populations, including second heart field (SHF)-derived regions. Importantly, no lineage-specific or strain-dependent biological comparisons are performed in this study; fluorescent labeling is used solely to visualize and track tissue motion. Acquisitions were performed at a variety of temporal windows and with variable time-lapse periods. The procedures developed therefore cope with a diversity of labelling strategies, temporal rates and embryonic periods of acquisition (Raiola et al., 2025).

To estimate tissue motion, we applied the Medical Image Registration Toolbox (MIRT) (Myronenko & Song, 2010), which calculates displacement tensors across time on raw images (Figure 1A) (Raiola et al., 2025; Myronenko & Song, 2010). This algorithm produces Tensor vectors (T) that describe the displacement of each image point over time (Figure 1A, orange grid). To validate these computational predictions, we compared them to manual tracking of >9000 genetically labelled cells derived from 9 embryos through consecutive frames (Raiola et al., 2025). The validation metric was based on the distance between predicted and experimentally measured positions. The results show that the predicted displacements closely match real cell movements, with an average error at the subcellular level, confirming that MIRT provides high-resolution and accurate quantification of tissue deformation during HT formation.

To generate a unified model of cardiac tissue motion, we registered each time-lapse sequence to the 3D+t Atlas of heart development (Esteban et al., 2022). First, we segmented the myocardial tissue in each specimen of the time-lapse datasets and produced a dense mesh of triangles that describes the tissue surface (Raiola et al., 2025). These meshes describe the dynamics of myocardial tissue motion and were named “Live-Shapes”. Second, we applied a machine learning-based staging system that uses morphometric features to align live images of different specimens over time, allowing the assignment of each specific specimen and stage to the corresponding Atlas geometry (Figure 1B) (Raiola et al., 2025). Third, we mapped specimens onto the Atlas to address shape variability between equally staged hearts (Figure 1B) (Raiola et al., 2025).

These steps allowed the mapping of parameters, such as strain and growth, from individual specimens onto the Atlas, providing their distribution and regional variability among specimens (Figure 1C, example of tissue growth mapping). Finally, using the tissue deformation data, we performed *in-silico* fate mapping by simulating the forward displacement of pseudo-cells placed in the CC domain. This analysis showed how different CC regions contribute to specific domains of the forming HT and how they deform during this process (Figure 1D).

### Extracting Deformation Patterns from the Continuous Description of Heart Tube Morphogenesis

To analyze tissue deformation patterns in the collection of Live-Shapes, we applied the continuous mechanics laws to the deformation of the mesh triangles (Figure 2A, (Raiola et al., 2025). Deformation was calculated between consecutive time points for 11 embryos, each with at least two shapes classified in successive Atlas stages. A subset of embryos was excluded because a substantial portion of the IFT was lost due to embryo drift, ensuring that all regions of the cardiac tissue are represented by the same number of embryos in the cumulative deformation analysis. The resulting pattern was then mapped onto the following stage shape, capturing the immediate deformation history. For each triangle, we computed local shape changes and applied smoothing (Raiola et al 2025) to estimate local tissue growth rate (J), tissue anisotropy (θ) and the magnitude and direction tissue deformation (ε) (Figure 2B). In addition, we introduced a new parameter; “Strain Agreement Index” (φ), which quantifies local coordination of strain directions. This metric distinguishes regions of coordinated deformation (ordered) from areas with locally discrepant deformation directions (chaotic). It would also identify borders between regions with coherent but discrepant strain directions (Figure 2B, MATERIALS AND METHODS - Strain Agreement Index). All deformation measurements are invariant to translation and rotation, making them robust to embryo drift during acquisition.

**Figure 2:**
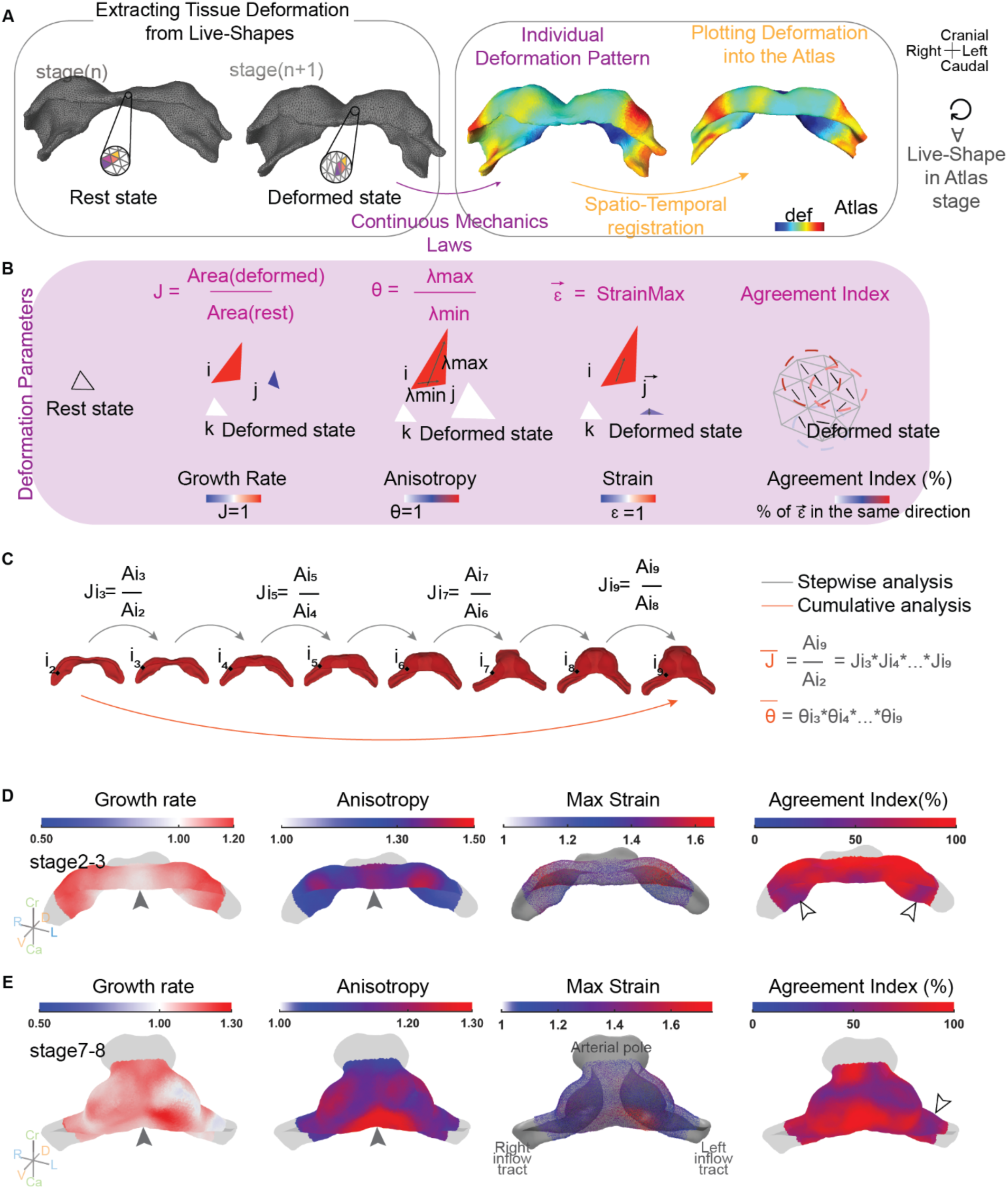
Deformation pattern analysis. A), Mesh transformation from the initial state to the deformed state. The triangles of the mesh are transformed according to the MIRT tensor from time ti to time ti+1. The deformation pattern is calculated by applying the principles of continuum mechanics between the initial and deformed states. This pattern is then visualized on the deformed shape. The spatio-temporal registration onto the Atlas enables the mapping of the deformation into it. B), Description of the deformation parameters between the initial and deformed states. For each triangle, three different possible deformations (a, b, c) are represented, color-coded according to the parameter value. The growth rate (J) is defined as the ratio of the area of the same triangle before and after deformation. Anisotropy (θ) is defined as the ratio between the maximum and minimum deformation magnitudes. The maximum strain (ε) is characterized by the vector of maximum deformation. The agreement index reports the local coherence in the direction of tissue deformation. C), Difference between stepwise deformation pattern analysis (grey arrows) and cumulative deformation pattern analysis (orange arrows). D-E), Stepwise deformation pattern analysis in caudal view. The color maps represent the growth rate, anisotropy, strain magnitude and direction, and finally, their agreement. Here, the deformation maps are shown for the transition from stage 2 to stage 3 and from stage 7 to stage 8. The full arrowhead indicates a zone of low growth rate and high anisotropy in the most ventral-medial region. The empty arrowhead indicates discontinuity in the agreement of the strain direction. Deformations values are the mean values. The number of specimens averaged at stages 2–3 is 5, while at stages 7–8 it is 2.

We then examined myocardial deformation pattern in two complementary ways: stepwise, by comparing deformation between consecutive stages; and cumulatively, by integrating the stepwise deformations through several stages (Figure 2C). To specifically focus on heart tube morphogenesis driven by tissue deformation, we excluded Atlas stage 1, given that between stages 1 and 2 substantial cell addition to the heart tube through myocardial differentiation takes place (Esteban et al., 2022).

### Stepwise Deformation Pattern Analysis

To investigate the spatial consistency of the deformation patterns among specimens, we mapped each deformation value from the Live-Shapes onto the corresponding Atlas geometry. For each stage group, we computed the spatial distributions of the mean deformation (μdef) (Figures S1-S3) and its variability (σdef) (Figure S4), with Figures S5–S6 showing the individual embryo contributions to these mean and std values. Importantly, while inter-embryo variability reflects natural biological diversity (Figure S4), it remained highly localized, primarily affecting deformation magnitude rather than spatial patterns, with anticorrelated growth/anisotropy zones consistently preserved across specimens (Figures S5-S6).

The stage2-to-stage3 and stage3-to-stage4 deformation analyses showed an inverse distribution of high growth and high anisotropy regions. High growth was associated with low anisotropy and high anisotropy appeared in regions with low growth. High anisotropy/low growth was mainly detected in a coherent region spanning the ventral midline region of the HT and extending laterally to the ventrocaudal region adjacent to the peripheral splanchnic mesoderm (see solid arrowheads in Figure 2D, and asterisks in Figure S1, Growth and Anisotropy columns). A second high anisotropy/low growth region appeared at the dorsal lips of the HT (see solid arrowheads in Figure S1, Growth and Anisotropy columns). The rest of the HT, including the IFTs, showed a largely isotropic growth (Figure S1). In stage 4, a left-biased increase in growth was detected (Figure S1, Growth column). Most specimens at this stage presented a coherent deformation direction, with strain oriented along the craniocaudal axis. Accordingly, they generally showed a high Agreement Index, indicating coordinated deformation of the myocardium. An exception to this is found at the lateral parts of the IFTs, where narrow bands of local strain disagreement were found separating the medial from the lateral parts of the IFTs (see empty arrowheads in the Agreement column). A closer observation of the strain alignment in these regions (see magnifications in the Max Strain column of Figure S1) shows that the low-agreement bands correspond to an abrupt transition between regions of coherent but discrepant strain directions. Whereas in the lateral-most regions of the IFTs strain aligns parallel to the main IFT axis, in their proximal region strain aligns craniocaudally, coherently with the rest of the HT (Figure S1, Max Strain and Agreement columns). These results indicate a strong compartmentalization of collective cell behavior during HT morphogenesis. This compartmentalization affects both the anisotropy/growth patterns and the directionality of tissue deformation. Furthermore, this analysis detected a strong anticorrelation between tissue growth and anisotropic deformation.

In stage 5 and stage 6, the medio-caudal myocardium continued to deform anisotropically along the craniocaudal direction, with a high rate of agreement, indicating coordinated deformation (Figure S2). At this stage, high growth rates are localized to the ventrolateral regions and extend into the inflow regions from the lateral regions of the forming ventricle (Figure S2). In general, high growth rates are mostly present in the caudal regions of the myocardium, except for some medio-lateral areas of the forming ventricle at stage 6 (Figure S2). This is accompanied by the appearance of regions of low agreement in the bilateral bulging areas of the forming ventricle (Figure S2). With some exceptions, growth and anisotropy continued to show anti-correlated patterns (Figure S2). The inflow regions again showed sharp boundaries between regions of high agreement with discrepant directions, indicating high compartmentalization of the deformation directions.

In stage 7 and stage 8, the caudal region of the myocardium showed reduced growth, whereas new high-growth areas appeared in the ventromedial region of the forming ventricle, extending bilaterally towards the cranial region around the arterial pole (Figure 2E, Figure S3). In all regions of the forming ventricle, growth correlated with a high index of agreement and, in general, with low anisotropy (Figure S3). The inflows continued showing frontiers of strong disagreement separating the distal inflow, predominantly showing a medio-lateral deformation direction, from more proximal regions, showing craniocaudal deformation directions (Figure S3). In stage 8, a left-side-specific increase in growth was observed in the ventral side of the ventricle.

In stage 9, the most prominent feature was the rightward rotation of the ventricle (Figure S3), a phenomenon related to heart looping (Le Garrec et al., 2017). This was accompanied by a sharp increase in growth and decrease in anisotropy at the left junction between the IFT and the ventricle, whereas the opposite took place on the right side (Figure S3). As in stage 8, higher growth was also observed in the ventral left side of the ventricle, suggesting its relationship to ventricle rotation.

An interesting observation throughout this study is the behavior of the myocardial rim in contact with the splanchnic mesoderm at the prospective dorsal pericardial wall. This border of the CC deforms drastically towards the cranial pole to form the rim of the arterial pole and the dorsal myocardial lips that fuse forming the mesocardium. This complex deformation is essential for the transformation of the CC into primitive HT. The dorsal and ventral views of the myocardium from stage 3 to stage 6 (Figures S1, S2) show that this rim is divided into two regions with different behaviors; a medial region that coincides with the closing dorsal lips and deforms towards the cranial pole, and bilateral distal regions, remaining at their original position. Interestingly, the medial aspects of this rim show very reduced growth or even contraction, whereas its lateral aspects show high growth rates. These results thus show again a strong compartmentalization of growth and deformation during HT formation.

### Cumulative Deformation Pattern Analysis

To determine the accumulated changes through several stages, we applied an additional strategy to analyze tissue deformation in chosen time windows (Raiola et al., 2025). The purpose was to recreate the continuous motion profile of the heart tissue during its transformation from CC to HT, exploiting the extracted kinetic information from individual live-imaging sequences. This strategy concatenates the motion profiles derived from the live images to fully cover from stage 2 to 9, allowing the cumulative deformation analysis (Raiola et al., 2025) (Figure 3A).

**Figure 3:**
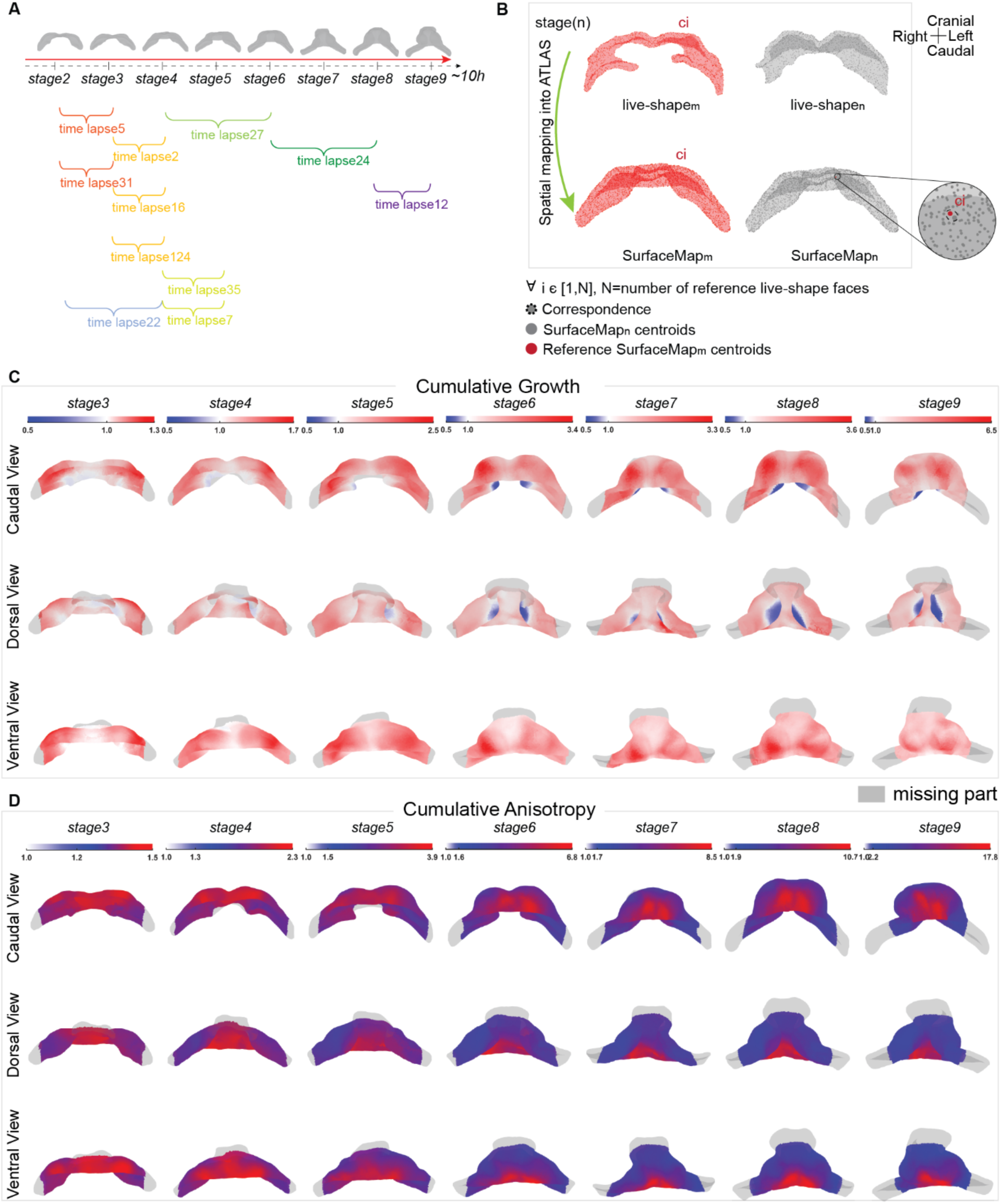
Cumulative Deformation Pattern Analysis. A), Concatenation of multiple time-lapses on a common timeline. Each time-lapse covers sub-windows of the Atlas temporal line. B), Schematic description of the concatenation pipeline. We fix a reference sample and for each equally staged sample, we select the centroids of the SurfaceMap closest to the reference SurfaceMap. C), Cumulative growth of HT. The color map refers to the mean growth rate of the tissue from stage 2. D), Cumulative anisotropy of HT. The color pattern indicates which zone of the myocardium accumulates more anisotropic deformation. The color map refers to the mean accumulated anisotropy of the tissue from stage 2. The color bars indicate the deformation magnitude. Grey zones correspond to the missing IFTs and arterial pole parts.

We used the projections of the Live-Shapes onto the Atlas, which we called “SurfaceMap” (Figure 3B), to establish correspondence between specimens and concatenate their motion profiles (Raiola et al., 2025). In this way, we could map the accumulated growth from stage 2 through the different stages. We found that the highest growth accumulated in the IFTs and at bilateral ventrolateral areas of the forming ventricle, whereas the midline and the arterial pole remained at low growth rates (Figure 3C). The two ventricular bilateral growth regions appeared as focal areas, with a clear single center from which growth rates decline concentrically (Figure 3C, ventral view). These high-growth foci progressively displace medially during development but always keeping bilaterality up to stage 9. The rim of the myocardium in contact with the prospective dorsal pericardium again showed two distinct regions; a bilateral distal region of relatively high growth in the IFTs, and the region of the dorsal closure, which shows low growth (Figure 3C). In contrast, anisotropic deformation accumulated along the ventral midline and the dorsal closure region of the myocardial rim, whereas it remained low in the expanding regions of the tissue (Figure 3D). These results confirm the strong compartmentalization of growth and deformation, with growth predominating in lateral regions, and deformation along the craniocaudal direction in medial regions

### In-silico Fate Map Analysis and Its Use in Determining Tissue Motion during Heart Tube Morphogenesis

Next, we used the concatenation strategy to create a deterministic *in-silico* fate map reproducing the motion profile of the myocardium during early morphogenesis. This strategy involved displacing the reference SurfaceMap nodes in accordance with the motion profile, generating a “Dynamic Atlas” as described in Raiola et al., 2025. In this Dynamic Atlas, every vertex of the model can be tracked from stage 2 to stage 9 (see Video S1). Given the correspondence between the model and the actual cell displacement (Raiola et al., 2025), the resulting *in-silico* fate map can be interpreted in terms of pseudo-cellular displacement within the myocardium. We next validated the Dynamic Atlas by developing the tool “fate map”, which allows to label any position/s and determine its fate *in-silico*. At the regional level, we compared the predicted regional growth in the Dynamic Atlas with that directly measured from the *ex-*vivo labeled cultured embryo. We performed experiments on *ex-vivo* cultured embryos using the TAT-Cre injection (Sendra et al., 2023) and dye labelling (Domínguez et al., 2012) to experimentally determine the fate of specific CC regions (Table S1). Using the TAT-Cre technique, we labelled a discrete group of cells in the medial CC and tracked the positions of the cell progeny after culturing the embryo for 20 hours (Figure 4A). An equivalent label generated in the stage 2 virtual model generated a labelled region in the stage 8 virtual model similar to that obtained in the experimental setting (Figure 4A, A’). Next, we labelled a group of embryonic cells with dye and examined the fate of the label after 15 hours (Figure 4B). We then labelled equivalent positions in the stage 2 virtual model and found that the fate of this region in the stage 6 virtual model was similar to that obtained in the experimental setting (Figure 4B, B’).

**Figure 4:**
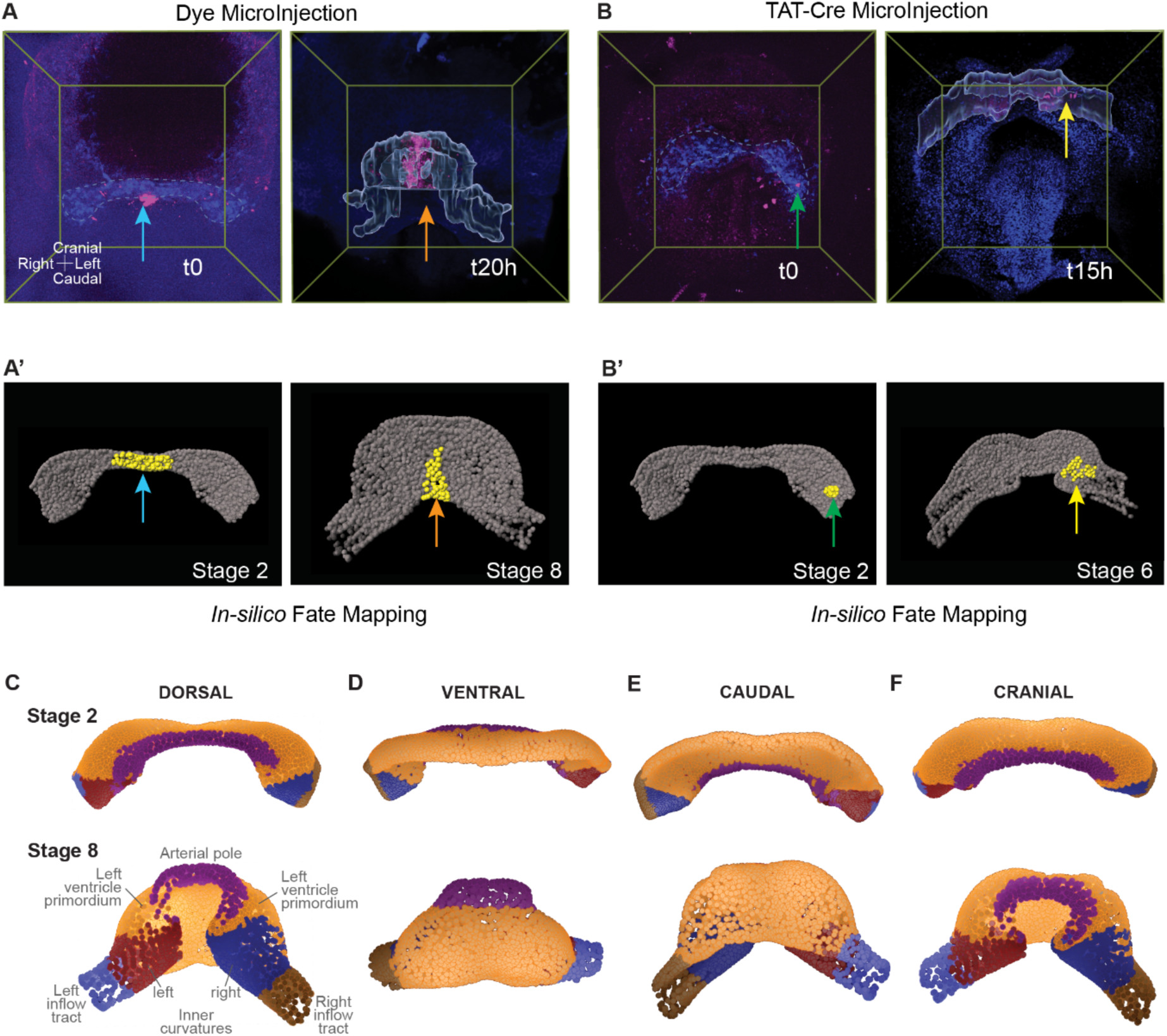
In-silico Fate Map. A), Labelling of a CC discrete region by dye injection (t0, left) and 3D reconstruction of its contribution to the HT after 15 hours of embryo culture (t15h, right). A’), labelling on stage 2 virtual model of an equivalent region to that experimentally labelled in A) (left) and the resulting labelled region in the stage 6 virtual model (right). B), Labelling of a cardiac crescent discrete region (t0, left) by TAT-Cre-induced recombination and 3D reconstruction of its contribution to the HT after 20 hours of embryo culture (t20, right). B’), labelling on stage 2 virtual model of an equivalent region to that experimentally labelled in A) (left) and the resulting labelled region in stage 8 virtual model (right). C-F), different views of the virtual model showing different regions of the HT in stage 8 and their primordia in stage 2.

Once determined that the virtual model approximates the behavior of myocardial cells during HT formation, we used it to map the primordia of the different regions of the HT observable at stage 8 (Figure 4C-F). We labelled the arterial pole, the left ventricle primordium (outer curvature), the ventricular side of the future inner curvature (dorsal side of the HT opposing the forming ventricle), and the IFTs. The comparison between the primordia and the end-stage structures indicates that the lateral-most parts of the CC grow and extensively migrate medially to generate the IFTs and the dorsal part of the HT (future inner curvature), whereas the primordia of the future ventricle (outer curvature) and the arterial pole mainly undergo deformation to achieve their final shape. Whereas both the arterial pole and the future ventricle reshape towards the midline, the arterial pole region contracts to a cylindrical shape, while the future ventricle bulges out in a barrel shape (Figure 4C-F).

Next, to understand how the ventricular barrel shape is produced, we used the virtual model to label groups of cells at regular spacing along the rims between the splanchnic mesoderm and the CC at stage 2 (Figure 5). These two rims define two lines, one between the CC and the juxta-cardiac field (Figure 5A; D2, as described in Esteban et al., 2022) and another between the CC and the SHF (Figure 5B; D1, as described in Esteban et al., 2022).

**Figure 5.**
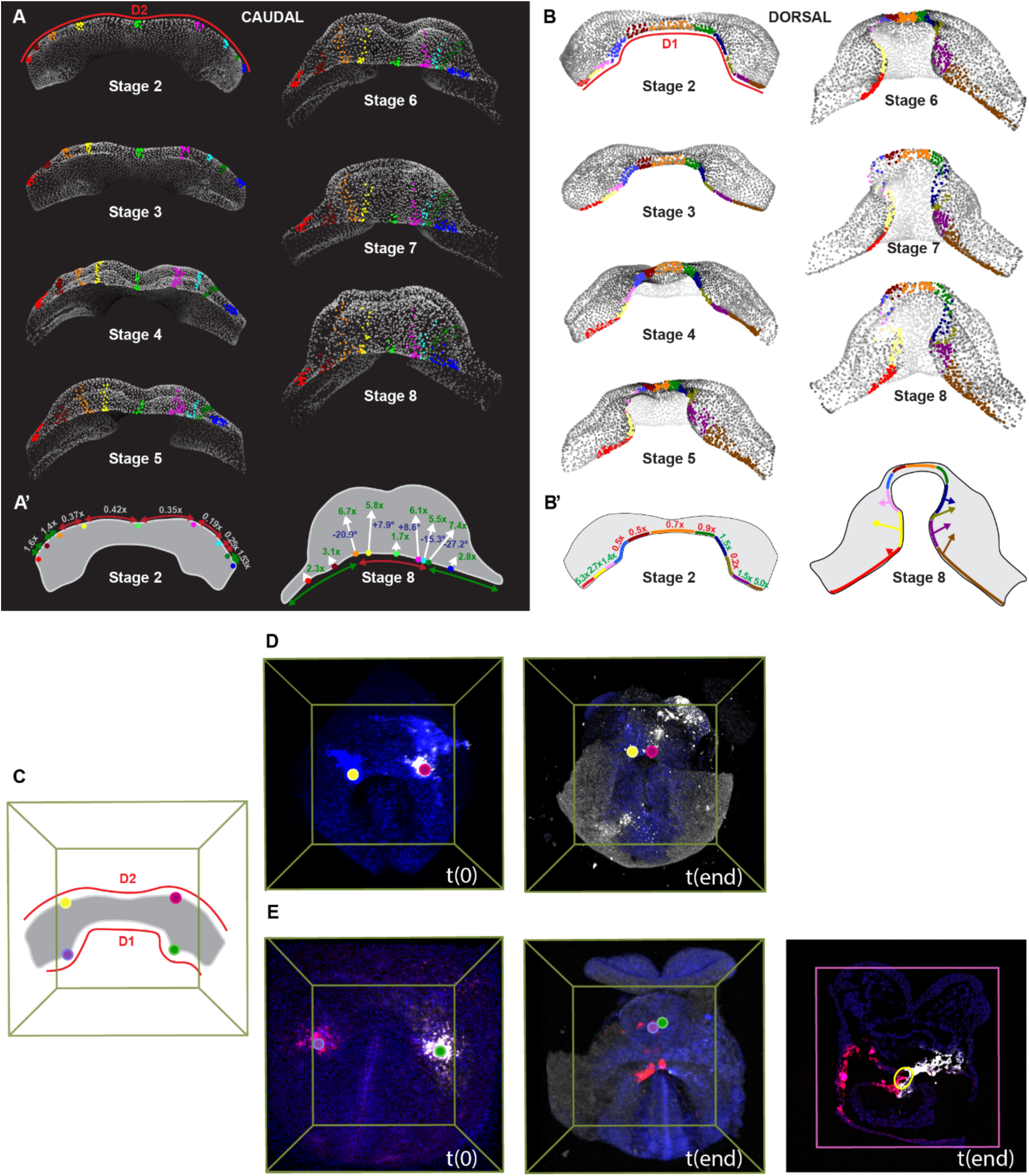
Fate maps of the cardiac crescent boundaries reveal tissue dynamics of heart tube formation. A), Clusters of pseudo-cells were labelled at regular spacing along the D2 line of the stage 2 CC model. The labeled clusters were then followed through stage 8. Caudal views are shown, which allow visualize the caudal side of the developing inflows and ventricle. A’) Left, representation of the degree of expansion (green) or contraction (red) of the different D2 segments defined between the labelled clusters at stage 2. Numbers indicate fold change from the initial to the final time points. Right, representation of the stage 8 view indicating the regions of D2 that contract (red) versus those that expand (green). Fold-change cranial ward expansion of each labelled cluster from their original craniocaudal extension is shown in green. The direction of the white arrows indicates the main direction of expansion. The angle of the main expansion direction with respect to the embryo mid-plane is indicated in blue. B), Contiguous segments of pseudo-cells were labelled at regular spacing along the D1 line of the stage 2 CC model. The labeled clusters were then followed through stage 8. Dorsal views are shown. B’) Left, representation of the labelled segments on stage 2 CC with indication of their fold-change expansion (green) or contraction (red). Right, Representation of the final extension at stage 9 of the D1 labelled segments. Arrows indicate the direction and extension of the expansion of the labelled pseudo-cells to colonize the dorsal parts of the HT. C), Schematic representation of the cardiac crescent at stage 2 indicating the four anchor points used for experimental validation: two along D2 (yellow and magenta dots) and two along D1 (purple and green dots). D) Experimental validation of D2 contraction. An embryo (e6_D2) was microinjected at the D2 anchor points (yellow and magenta) and imaged by multiphoton microscopy at t0 (left) and after 10 hours (right). The Euclidean distance between the two labeled anchor points was measured at both time points (right image). E), Experimental validation of D1 contraction. An embryo (e1_D1) was microinjected at the D1 anchor points (purple and green) and imaged at t0 (left) and after 10 hours (centre). Given the difficulty of imaging the arterial pole at high resolution by whole-mount microscopy, cryosections were performed (right, pink frame) to measure the geodesic distance between anchor points in a coronal plane (arterial pole in yellow). Multiphoton images were acquired at a voxel size of 0.57 × 0.57 × 2.5–6.0 µm. Cryosection images were acquired at a pixel size of 0.65 × 0.65 × 0.042 µm.

This study shows that along D2 only the lateral-most sections expand towards the midline to generate the sections of D2 underlying the IFTs at stage 8 (Figure 5A, A’). In contrast all D2 sections underlying the ventricle at stage 8 undergo a strong compression from their initial length at stage 2 (Figure 5A, A’). Whereas the labels that end up positioned within the IFTs show limited cranial expansion, those underlying the ventricle extensively expand cranially and colonize the caudal part of the ventricle (Figure 5A, A’). At stage 8, the future left ventricle shows two hemi bulges with an indentation between them corresponding to the midline. The strongest lateral compression and cranial expansion are found for bilateral groups of cells close to the midline of these two hemi bulges, while the midline label expands only moderately towards the ventricular wall (Figure 5A, A’). Interestingly, the cranial extensions of these bilateral cell groups open laterally as they extend cranially, generating a fan-like label (Figure 5A, A’). These observations suggest that the lateral constriction and cranial expansion of cells close to D2 contribute to generating the barrel shape of the ventricle.

The same study on D1 shows that the region fated to the arterial pole compresses to form the cylindrical shape of this structure, whereas the lateral aspects of D1 expand drastically to form the cranial border of the IFTs and the dorsal borders of the ventricular region, which will later fuse to form the mesocardium (Figure 5B, B’). In this future mesocardial region, cells originally positioned close to D1 expand to populate the inner curvature of the ventricular region (Figure 5B, B’). These deformations ensure that the HT forms by generating the inner curvature regions and allowing its dorsal closure. At the same time, the constriction imposed by the formation of the arterial pole limits the expansion of the ventricle at its cranial end, thereby contributing to its barrel shape.

To experimentally validate the tissue dynamics predicted by the virtual model, we identified four anchor points, two along D1 and two along D2, to track changes in these boundaries during heart tube formation (Figure 5C). For D2, three embryos were microinjected at the defined anchor points, imaged around stage 2, and re-imaged after 10–14 hours by multiphoton microscopy (Figure 5D). The Euclidean distance between the two D2 anchor points was computed at t0 and tend. In all three embryos, the anchor points converged over time, with the D2 segment retaining on average 0.27 ± 0.14 of its initial length (range: 0.13–0.40), reflecting a substantial contraction whose magnitude varied depending on the developmental stage and exact position of microinjection. Consistent with these measurements, the model predicted a contraction of approximately 0.38 between the yellow and magenta reference points (Figure 5A’), in agreement with the experimentally observed deformation at the corresponding locations in Figure 5D-E. For D1, three additional embryos were microinjected in the same manner and the in-plane geodesic distance between anchor points was measured at t0 and after 10–14 hours (Figure 5E). Given the difficulty of imaging the arterial pole at high resolution by whole-mount microscopy, cryosections were used. The D1 segment defining the arterial pole similarly underwent compression, retaining on average 0.50 ± 0.22 of its initial length (range: 0.23–0.77). In all cases, the reduction in both D1 and D2 distances over time is consistent with the contraction dynamics predicted by the barrel model (Table S2).

Finally, to understand the shape changes taking place during the formation of the primitive left ventricle, we labelled longitudinal rectangles in the ventricle of the stage-8 model and studied the evolution of their shape from stage 2 to stage 8 (Figure 6A). The observations indicate that the whole ventricular primordium undergoes extensive reshaping with progressive constriction of the tissue towards the midline in a craniocaudal gradient. The movement resembles the closing of an open fan with the vertex at the cranial side (Figure 6B). Through this movement, lateral tissue is recruited towards the midline and extends cranial ward. An exception to this general reshaping is the mid-central region of the ventricle, where bilateral deformation takes place in mirror-image directions with an equatorial axis of symmetry. This deformation pattern correlates with the localized fan-like distribution of the descendant of D2 (Figure 5A, A’, indicated by asterisks in Figure 6). These observations therefore match the deformations observed for labels in D1 and D2 (Figure 5). Together, these results show that the ventricle primordium is narrower at the cranial end of the CC and extends progressively more laterally towards the caudal end of the CC. Ventricle morphogenesis thus involves tissue constriction by two “belts” of tissue that support the barrel shape of the ventricle. The cranial belt constitutes the arterial pole and the caudal belt coincides with the myocardial region abutting the juxta-cardiac field. This caudal belt undergoes a much more extensive constriction than the cranial one, related to the recruitment of very lateral aspects of the CC. In contrast, at the cranial aspect of the CC, this constriction is less strong and ventricle formation does not involve the recruitment of the lateral-most CC regions.

**Figure 6.**
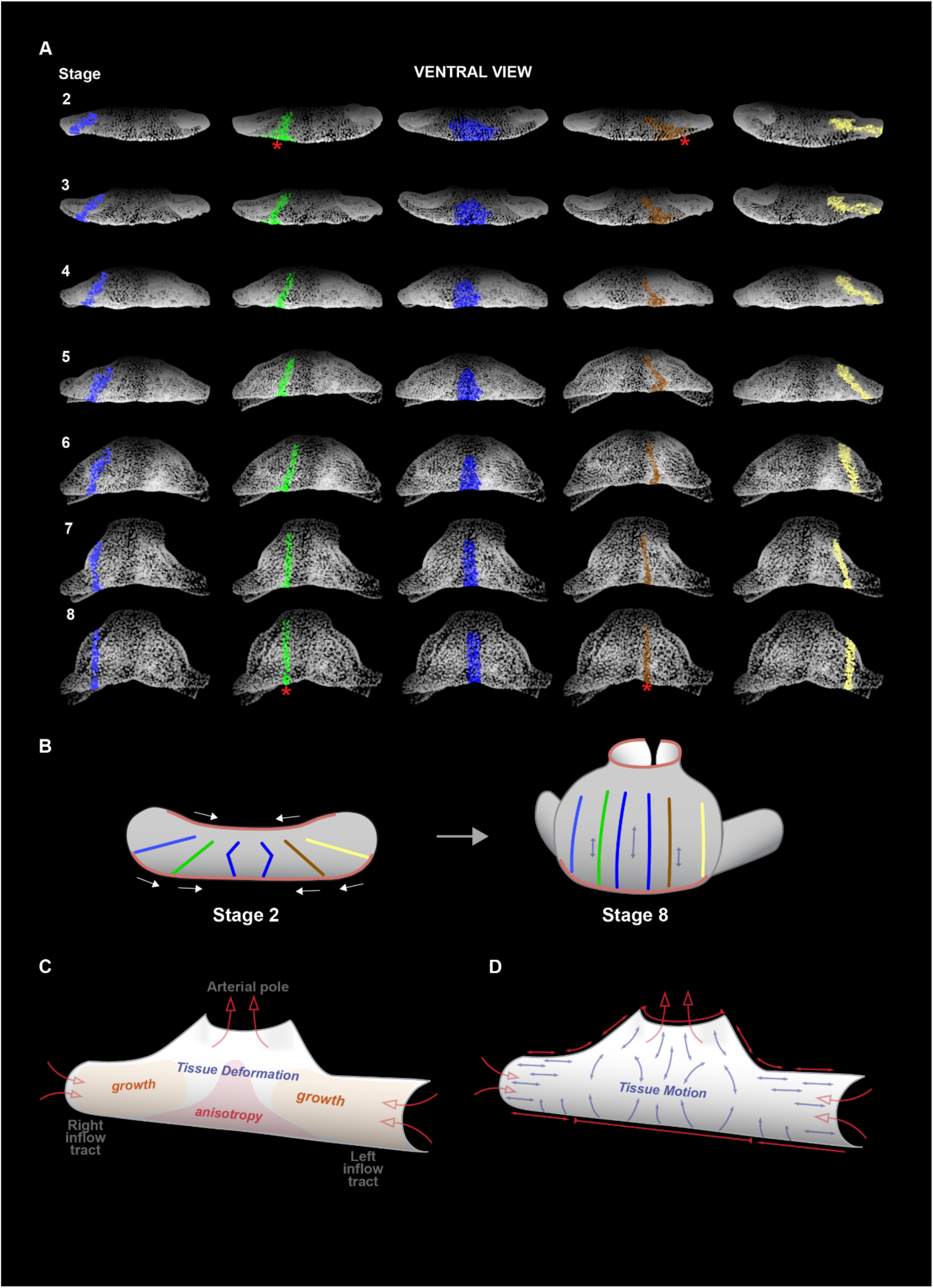
Fate map of the ventricle primordium shows the tissue dynamics underlying ventricle formation. A), longitudinal rectangular groups of pseudo-cells were regularly labeled in the ventricle of the stage 8 model. The labelled regions were then tracked back through stage 2. B), Schematic representation of regional deformations at stage 2 and stage 8. C) Representation of the main growth and anisotropy areas in the myocardium. D) Representation of the main directions of tissue motion during heart tube formation.

## DISCUSSION

Here, we applied a previously established image-based pipeline (Raiola et al., 2025) to extract and compare myocardial deformation during HT morphogenesis in the mouse embryo. Our analysis comprises 16 specimens that have been staged into 9 different canonical shapes at nominal ages ranging from E7.75 to E8.25 (approximately 12h). The canonical shapes provide a spatiotemporal reference for comparing and compiling the motion and deformation of multiple specimens.

The knowledge of the deformation patterns extracted from the videos allowed the cumulative evaluation of the most important transformations of the myocardium, measuring their intensity and direction, and determining their timing. The maps reveal a strong compartmentalization of growth rates and anisotropic deformation. Growth is generally high in the lateral regions of the forming ventricle, with a bias towards its caudal parts. In contrast, the medial stripe of the CC, later of the forming tube, does not appreciably grow but shows high anisotropic deformation with stretching mainly oriented in the rostro-caudal direction. Similar mutual exclusion of the high-anisotropic areas and the growing bilateral caudal regions is clearly maintained until stage 6. This compartmentalization generates clear boundaries between regions that show coordinated stretch directions and those that do not and between regions showing different stretching directions (Figure 6C). These boundaries are particularly pronounced in the inflow region, where differences in stretching directions become particularly evident (Figure 6D). In the IFT, boundary regions suggest two forces that act in different directions: one, stretching the IFT cells in the latero-medial direction, and another promoting the elongation of the tube in a craniocaudal direction. On the dorsal side of the forming tube, the rim that connects the myocardium to the splanchnic mesoderm (D1) also shows a high mediolateral compartmentalization, with its medial part undergoing little growth or even compression at several stages, whereas its lateral parts grow and stretch toward the midline (Figure 6D). This originates a progressive cranial looping of the medial aspects of this rim, which form the arterial pole and the closing mesocardium, whereas the lateral-most parts remain as IFTs.

To directly test whether the compression dynamics predicted by the virtual model occur in vivo, we performed microinjection experiments tracking anchor points along D1 and D2 over 10-14 hours (Figure 5C–E, Table S2). In all six embryos analyzed, anchor points converged over time along both boundaries, representing to our knowledge the first direct in vivo evidence that the medial sections of D1 and D2 undergo tissue contraction during heart tube formation. This observation indirectly implies the expansion of their lateral sections, provided both D1 and D2 expand during development, with D1 showing a much stronger expansion. The variability observed across embryos reflects the inherent challenges of this experimental approach: the exact position of the anchor points could not be perfectly standardized, embryos were not always captured at exactly the same developmental stage at t0 and tfinal, and high-resolution geodesic staging was deliberately avoided to preserve embryo viability. Nonetheless, the consistent direction of shrinking among specimens supports barrel model predictions.

Given that tissue compression inhibits proliferation through the activation of the Hippo pathway (Von Gise et al., 2012), we propose that one of the factors contributing to the lower growth of the medial regions might be tissue compression. Alternatively, or in addition, it is possible that forces arise from the splanchnic mesoderm, which is highly proliferative (de Boer et al., 2012) and could contribute to the anisotropic deformation of the mesocardial rim of the myocardium.

Based on these findings, we propose a model for mouse primitive HT morphogenesis in which areas of growth are predominantly lateral in the CC and later become ventrocaudal during the formation of the ventricle. Given that the lateral extension of the heart remains stable during HT formation (Esteban et al., 2022), we propose that the growth of these lateral regions causes the medial convergence of the tissue. This convergence is driven by two belts that constrain the CC towards the midline, a cranial one that coincides with the arterial pole and a caudal one at the boundary between the ventricle primordium and the juxta-cardiac field (Figure 6D). In between these two belt-like constraints, the ventricle adopts a bulging barrel shape (Figure 6B). Surprisingly, ventricle morphogenesis at this stage does not involve localized areas of growth but rather extensive remodeling to recruit lateral tissue and relocate it along the craniocaudal direction (Figure 6B). The model thus proposes complementary regions in which growth and deformation are anticorrelated. Concomitant with this recruitment to medial regions, HT tissue is deformed in the craniocaudal directions, so that the cranial and caudal belts move apart from each other and the HT adopts its longitudinal disposition (Figure 6B).

Between stage 7 and stage 9, we observed a difference in growth rate between the two sides of the forming ventricle, consistent with de Boer et al., 2012. This bias could be the first input for the looping process. Increased growth in the left region, as observed in the chicken model, could influence the direction of cardiac looping and result in a rightward C-shape (Kidokoro et al., 2008). As a consequence of the slight bending, the tissue could undergo compression on the right side. Alternatively, the observed bias could derive from differential growth of myocardial cells during looping (Ebrahimi et al., 2022; Shi et al., 2014). In addition, differential L-R proliferation in the dorsal pericardial wall may also influence heart looping by influencing the dorsal myocardial rim (Desgrange et al., 2020).

By concatenating the motion profiles of multiple specimens, we present the first in silico fate map, which enables virtual analyses of the dynamics of HT formation. The resulting Dynamic Atlas is a deterministic, descriptive representation of tissue morphogenesis, obtained by integrating individually derived motion fields within a common reference frame. This baseline is essential for studying heart defects, as deviations from normal tissue motion and deformation patterns can reveal developmental defects like altered growth or wrong motion paths. Due to its deterministic nature, the framework does not allow predictions of system responses to perturbations, but it provides a robust baseline for characterizing HT tissue dynamics. This approach reveals strong anisotropic deformation of the tissue, particularly in the ventricle and the dorsal myocardial boundary, where cumulative deformation analysis indicates the greatest anisotropy. Although the accuracy of the kinetic motion captured by the fate map has not been directly validated, we believe it can reliably track cardiomyocyte movement in these regions, as confirmed by comparison with the video-microscopy dataset (Raiola et al., 2025). We propose the Dynamic Atlas and its associated tool, the “fate map,” as a valuable approach for exploring HT morphogenesis.

In conclusion, by applying a previously established workflow (Raiola et al., 2025), we provide detailed kinetics of myocardial motion and the tissue deformations underlying early mammalian heart development. The fate map is proposed as an innovative tool to shed light on regionalized cell coordination, generating new questions about the biological and genetic factors that regulate the cardiac development process.

## MATERIALS AND METHODS

### Mouse Strains

Animals were handled in accordance with CNIC Ethics Committee, Spanish laws and the EU Directive 2010/63/EU for the use of animals in research. All mouse experiments were approved by the CNIC and Universidad Autónoma de Madrid Committees for “Ética y Bienestar Animal” and the area of “Protección Animal” of the Community of Madrid with reference PROEX 220/15. For this study, mice were maintained on mixed C57Bl/6 or CD1 background. We used the following mouse lines, which were genotyped by PCR following the original study protocols. Male and female mice of more than 8 weeks of age were used for mating.

### Embryo Culture and Live Imaging

Live imaging procedures were performed following the protocol described by Sendra et al. (2022). Mouse embryos (E6.5–E7.5) were collected and dissected in dissection medium consisting of DMEM supplemented with 10% fetal bovine serum, 25 mM HEPES–NaOH (pH 7.2), and penicillin–streptomycin (50 µg/ml each). Embryos were cultured in a medium composed of 50% Janvier Labs Rat Serum (Sprague Dawley RjHan SD, male only) and 50% DMEM FluoroBrite (Thermo Fisher Scientific, A1896701) at 37°C under 7% CO₂. Two-photon live imaging was performed on a Zeiss LSM780 microscope equipped with a 20× objective (NA = 1) and a MaiTai laser tuned to 980 nm for two-channel acquisition. Fluorescence signals were detected using non-descanned detectors with cyan-yellow (BP450–500/BP520–560), green-red (BP500–520/BP570–610), and yellow-red (BP520–560/BP645–710) filter sets. Image acquisition was controlled using Zen software (Zeiss) with an output power of 250 mW, a pixel dwell time of 14.8 µs, line averaging of two, and an image size of 1024 × 1024 pixels.

### Cell Tracking with TAT–Cre Microinjection and Dye Microinjection

Embryos at stages E6.5-E7.5 were dissected and cultured as described above using the same dissection and culture media. For microinjection experiments, embryos were maintained in a hypoxic chamber incubator at 37°C with 5% O₂ and 7% CO₂. Embryos were microinjected with TAT-Cre recombinase or fluorescent dyes as previously described (Sendra et al., 2023). Microinjection needles were prepared with a 2µm gauge and inserted into the anterior side of the embryo until penetrating the endodermal layer, using specified pressure conditions. The embryos were handled and positioned carefully, ensuring that the anterior and posterior sides were oriented accordingly during the procedure to achieve successful microinjections.

We utilised mouse embryos that carried both the reporter genes ROSA26CAG–TdTomato (R26RtdTomato) and ROSA26CAG–EGFP (R26REGFP) in transheterozygosis. We fixed, imaged, and annotated fluorescent cells within anatomical regions. “Clusters” were defined as groups of cells (either Tomato or GFP) originating from a single TAT-Cre injection.

For dye microinjections, mouse embryos were labeled by injection of a lipophilic carbocyanine, DiI (1,1’-dioctadecyl-3,3,3’,3’-tetramethylindocarbocyanine perchlorate) and DiD (1,1’-dioctadecyl-3,3,3’,3’-tetramethylindodicarbocyanine), as described previously (Domínguez et al., 2012). Subsequently, embryos were incubated in increasing sucrose concentrations (10%, 20%, and 30%) for tissue cryoprotection, embedded, and cryosectioned using a cryostat. Images were acquired by confocal microscopy.

### Genetic cell labelling

To visualize cardiac tissue and achieve fluorescent contrast for motion estimation and cell tracking, embryos expressed fluorescent reporters under the control of several commonly used cardiac or mesodermal drivers. These included Nkx2.5-GFP and Nkx2.5-Cre lines to label myocardial tissue, Mesp1-Cre to label mesodermal derivatives, and Islet1-Cre to label a population enriched in second heart field progenitors. We note that Islet1-Cre activity is not restricted to the second heart field and includes additional cardiac progenitor populations, including subsets of left ventricular precursors. However, this does not affect the analyses presented here, as lineage identity is not used for biological interpretation.

Fluorescent labelling was used solely to create sufficient signal contrast to follow tissue motion or to identify individual cells for validation of the motion estimation algorithm. The computational framework does not rely on tracking specific cell types, nor does it assume lineage specificity. Instead, fluorescence serves as a generic marker of tissue continuity and motion, allowing validation of the image-based registration independently of the genetic driver used.

For sparse cell labelling, embryos carried a ubiquitously expressed CreERT2 allele (Polr2a-CreERT2, RERT) in combination with Rosa26 reporter alleles. Recombination was induced by low-dose tamoxifen administration to pregnant females, resulting in stochastic and sparse labelling of cells. Tamoxifen was prepared by dissolving 4-hydroxytamoxifen (Sigma) at 1 mg/ml in corn oil with ethanol as an intermediate solvent, followed by sonication. A single 0,2 mg intraperitoneal injection was administered at E6.5. Sparsely labelled cells are used exclusively as fiducial markers to provide ground-truth trajectories for validating the accuracy of the deformation fields computed from image registration. Labelling efficiency and spatial distribution were therefore assessed only to ensure sufficient numbers of isolated cells for validation purposes, rather than to achieve comprehensive or lineage-restricted coverage.

### Computational Workflow

The computational workflow is detailed in Raiola et al. (2025). Here, we describe only the additional analysis introduced in this study that was not included in the methodological paper.

### Strain Agreement Index

We defined the stretch direction as the eigenvector related to the maximum eigenvalue. To better understand the stretching directions at the tissue level, we introduced an additional parameter called the Strain Agreement Index (φ). This parameter identified regions of the heart that stretched in the same direction, discriminating them from those with chaotic or discordant directions. φ was calculated as the percentage of directions that were in concordance in a certain neighbourhood. The choice of the neighbourhood size, corresponding to 6-7 cells (approximately 20 pixels), was defined experimentally to achieve a balance between local resolution and robustness. This scale was small enough to allow the tissue to be computationally flattened, avoiding artefacts from folded regions, while larger neighborhoods would have reduced local sensitivity. Conversely, smaller neighborhoods would have produced fragmented, salt-and-pepper patterns lacking generalization.

We evaluated the φ index for each face considering the stretching in this neighbourhood. As a first step, we projected the selected stretch vectors onto a plane identified by the centroids of the selected faces, using the *fitPlane* and *projLineOnPlane* functions of the geom3d library (http://github.com/mattools/matGeom/). In the second step, we calculated the angle formed by each stretch vector with the remaining vectors. We then evaluated the Agreement Index as the percentage of vectors whose angle diverged from ±10°. The percentage value, ranging between 0 and 100%, was converted into a color code and plotted for each mesh.

## Acknowledgements

We thank members of the Torres group for inspiring discussions and advice. We thank members of the Microscopy and Dynamic Imaging, Transgenesis, and Animal Facility CNIC units for excellent support. M.R. was a recipient of a Marie Skłodowska-Curie doctoral contract from H2020-MSCA-ITN-2016-722427. This work was funded by grants PGC2018-096486-B-I00 and PID2022-140058NB-C31 from the Agencia Estatal de Investigación to M.T.; Comunidad de Madrid grant P2022/BMD-7245 CARDIOBOOST-CM to M.T.; European Research Council AdG ref. 101142005 to M.T. The CNIC Unit of Microscopy and Dynamic Imaging is supported by FEDER ‘Una manera de hacer Europa’ (ReDIB ICTS infrastructure TRIMA@CNIC, MCIN). The CNIC is supported by the Instituto de Salud Carlos III (ISCIII), the Ministerio de Ciencia, Innovación y Universidades (MICIU) and the Pro CNIC Foundation, and is a Severo Ochoa Center of Excellence (grant CEX2020-001041-S funded by MICIU/AEI/10.13039/501100011033).

## Author contributions

Conceptualization: M.R., M.T.; Methodology: M.S., J.N.D, M.R.; Software: M.R.; Formal analysis: M.R., J.N.D.; Investigation: M.S., J.N.D., M.T.; Data curation: M.R., M.T.; Writing - original draft: M.R.; Writing - review & editing: M.R., M.T.; Supervision: M.T.; Project administration: M.T.; Funding acquisition: M.T.

## SOFTWARE

We recommend opening the 3D mesh with the open-source software ‘MeshLab’ (https://www.meshlab.net/). In ParaView (https://www.paraview.org), you can plot the MaxStrain vectors. Once the .vtk file is imported, the vectors can be visualized using the Glyph representation. It is recommended to display the vectors as lines for better visual interpretation.

To view and use the *in-silico* Fate Map, the .ims data needs to be imported into Imaris Viewer (https://imaris.oxinst.com/imaris-viewer), the free version of the Imaris visualization software.

## SUPPLEMENTAL ITEMS

**Figure S1:**
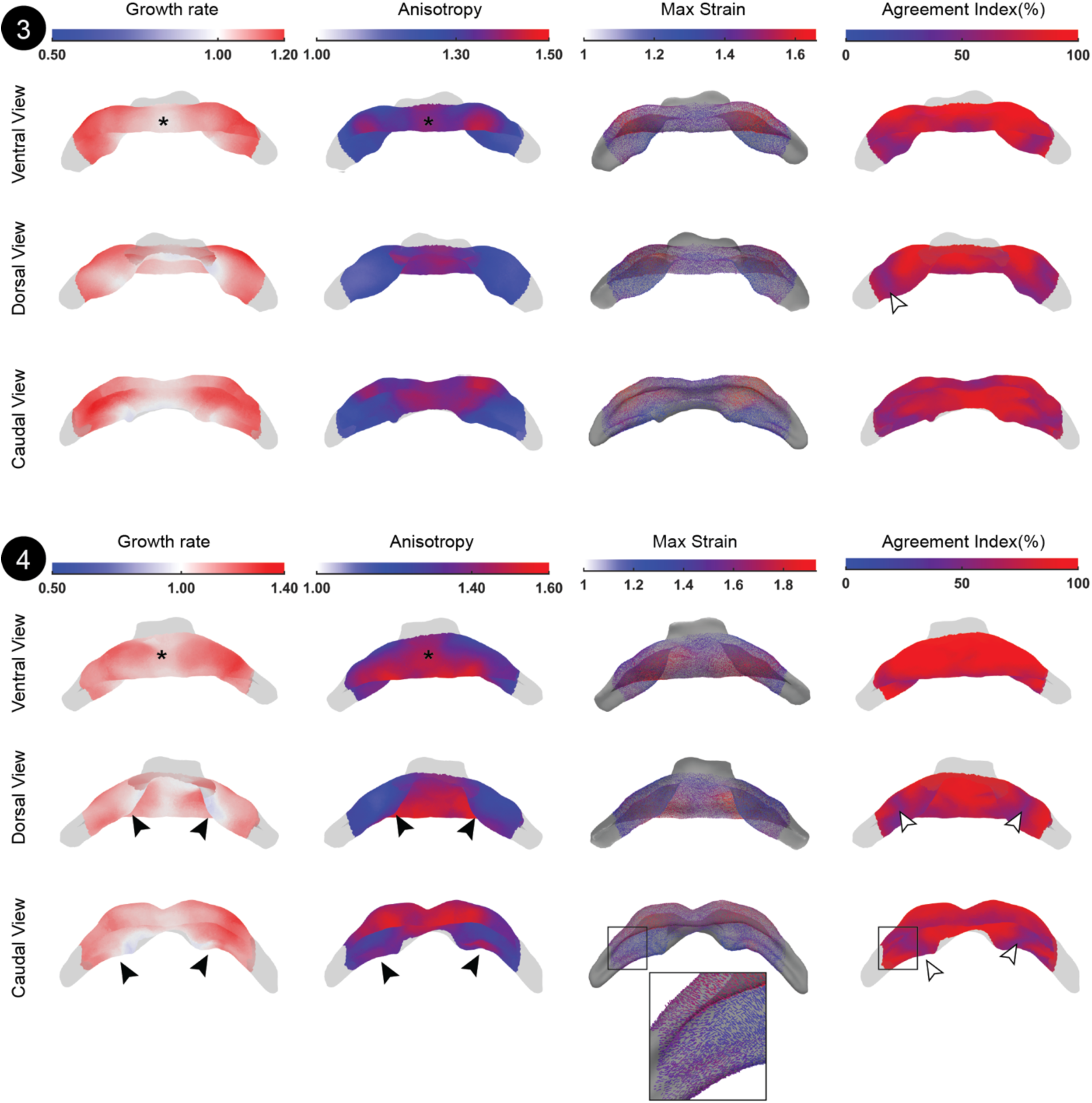
Stepwise Deformation Pattern Analysis. Deformation analysis for stage 3 and stage 4. The number of specimens averaged at stage 2 is 5, while at stage 4 it is 6.

**Figure S2:**
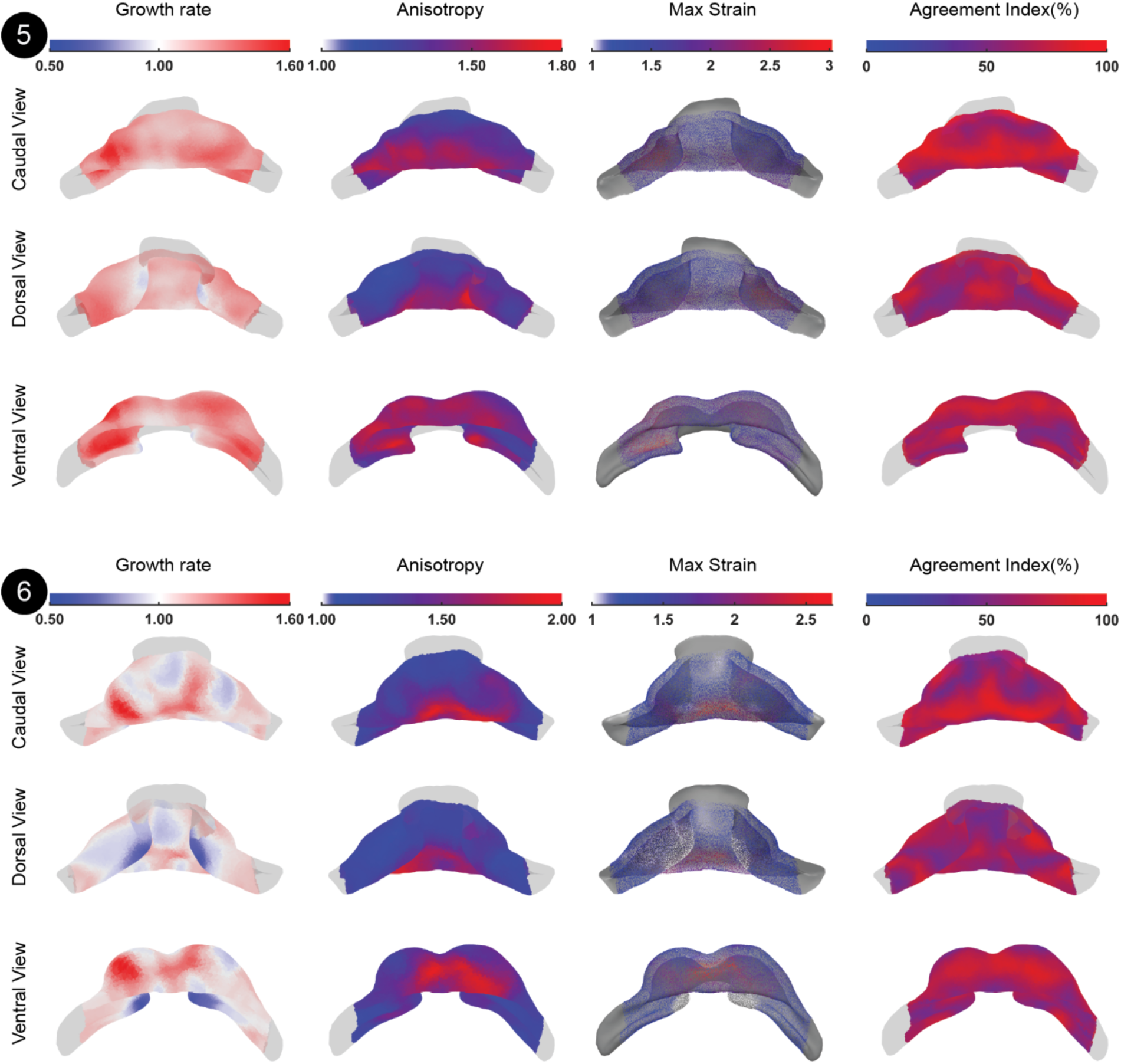
Stepwise Deformation Pattern Analysis. Deformation analysis for stage 5 and stage 6. The number of specimens averaged at stage 5 is 2, while at stage 6 it is 2.

**Figure S3:**
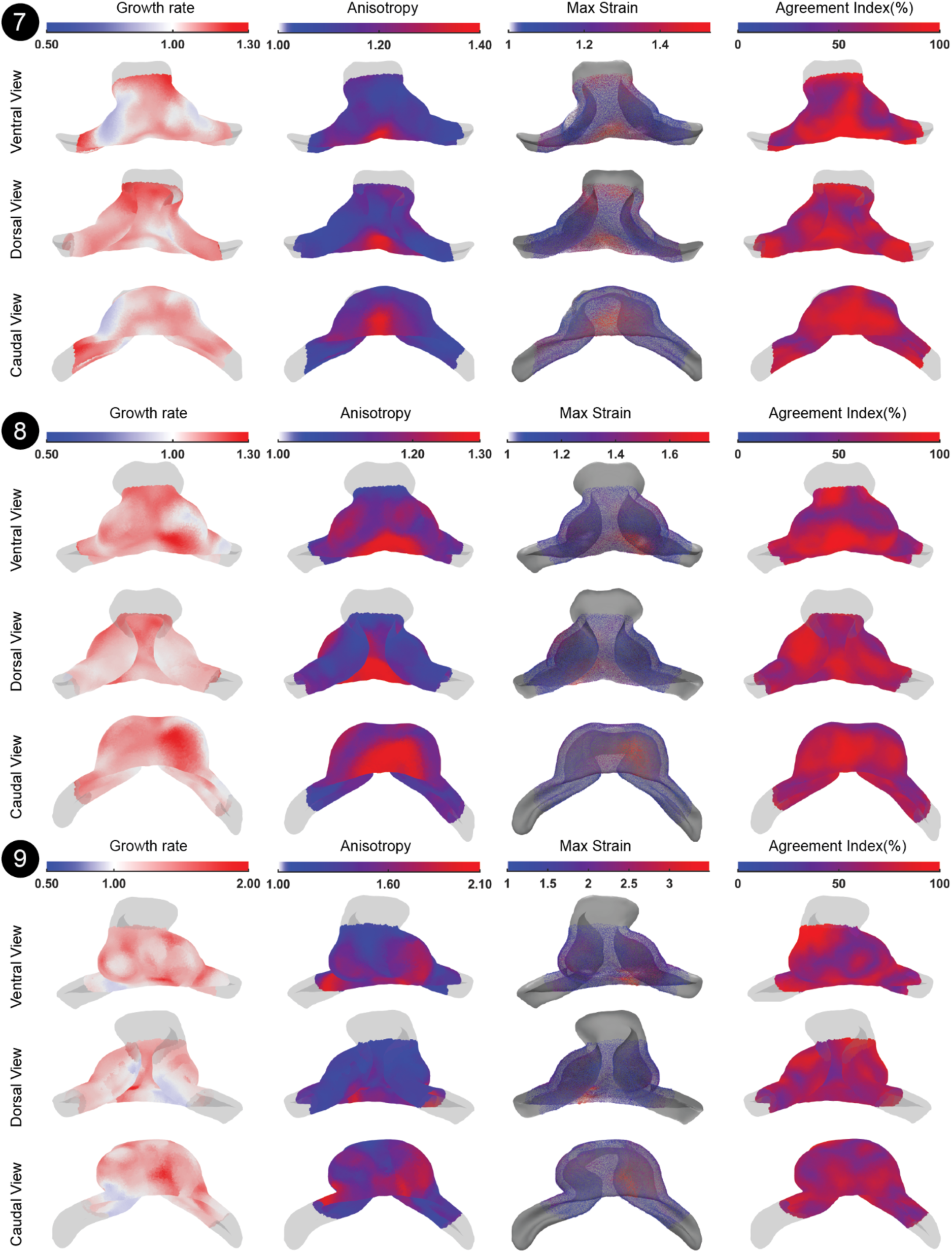
Stepwise Deformation Pattern Analysis. Deformation analysis for stage 7, stage 8 and stage 9. The number of specimens averaged at stage 7 is 1, at stage 8 is 2, and at stage 9 is 1.

**Figure S4:**
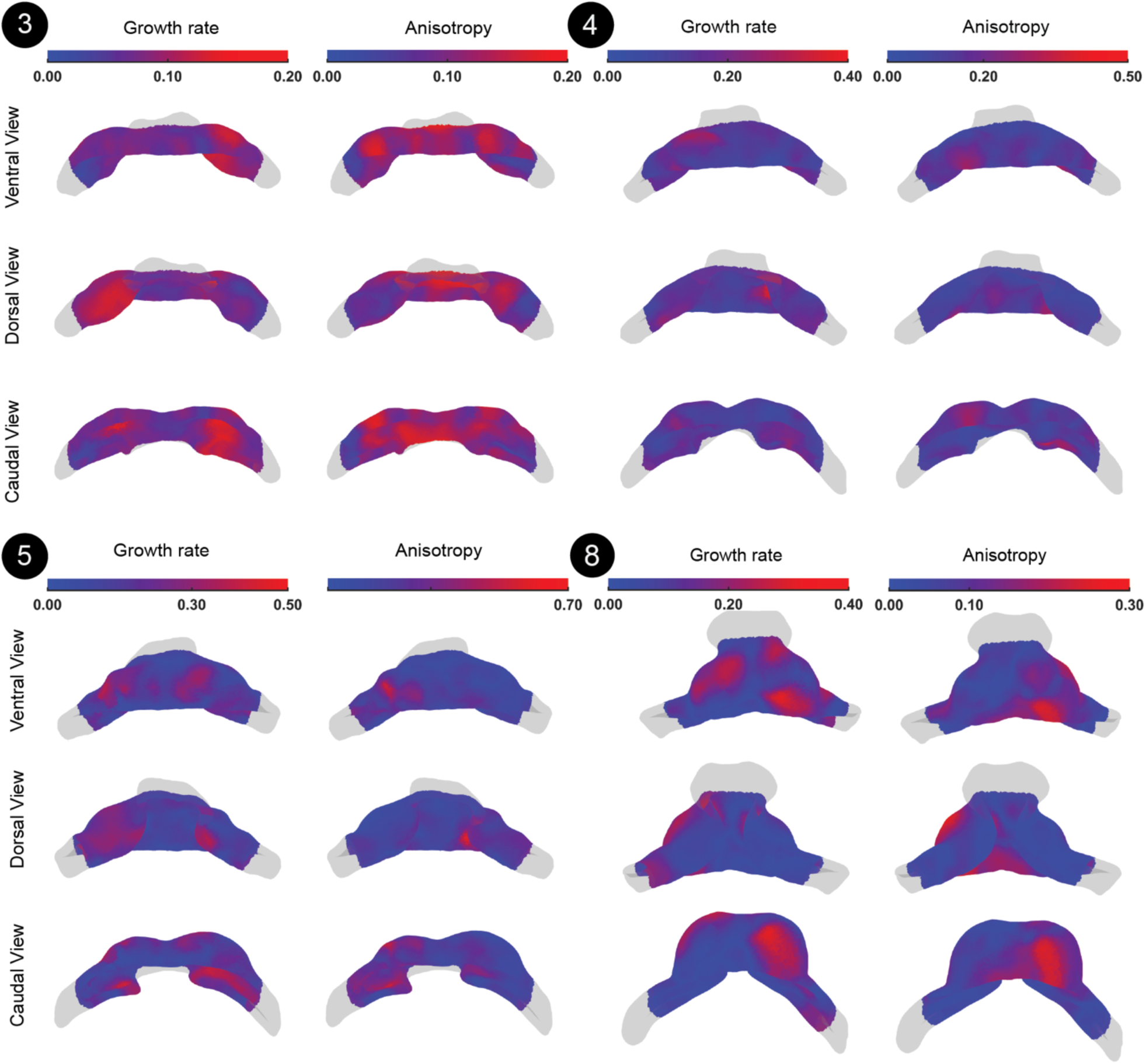
Standard deviation of the growth rate and the anisotropy patterns. Colormaps indicate the spatial distribution of the Standard Deviations for the growth rate and anisotropy magnitudes shown from Ventral, Dorsal and Caudal views. Stages 6,7,9 have only one staged Live-Shape and therefore variability could not be measured.

**Figure S5.**
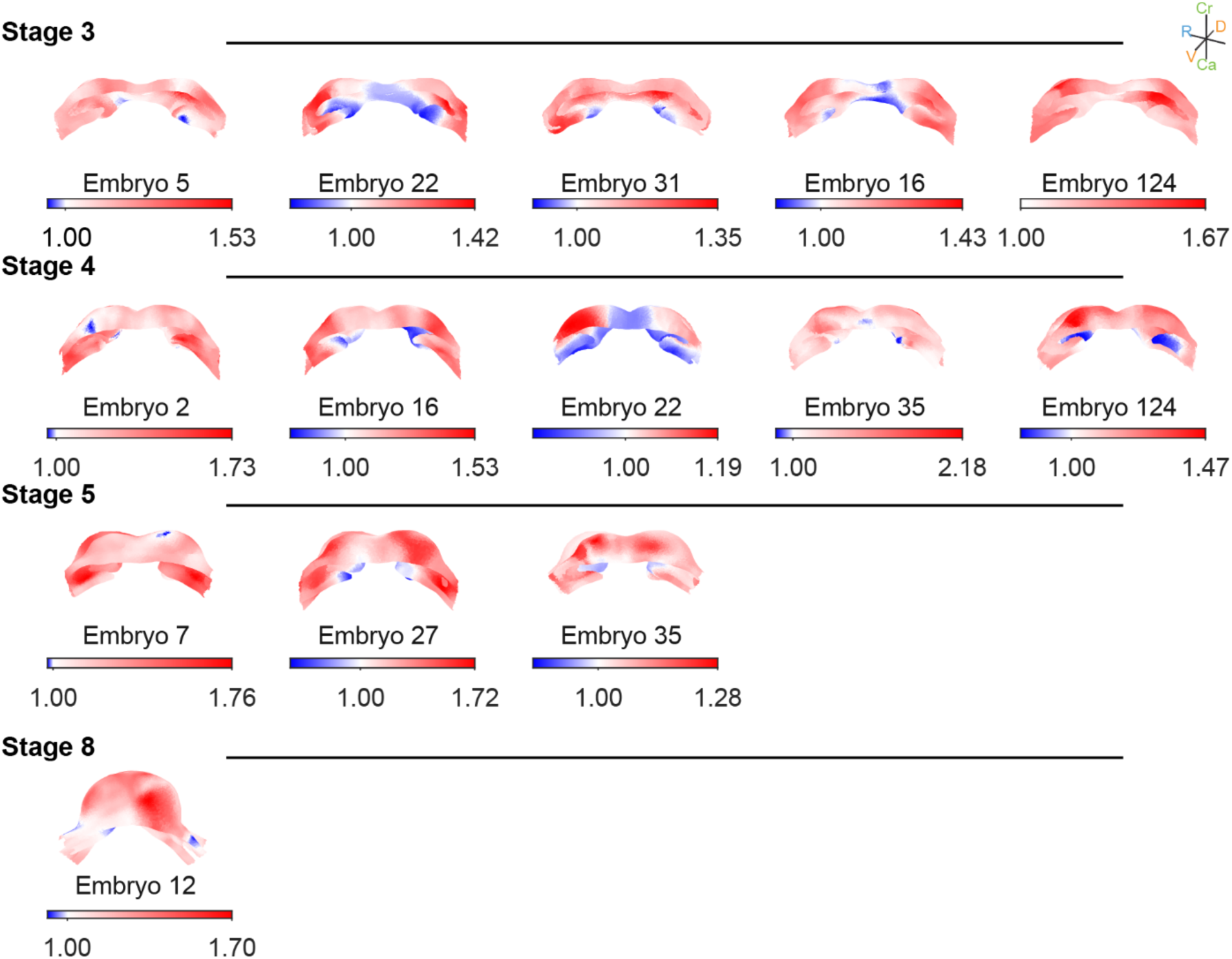
Mean growth rate derived from stepwise deformations between consecutive developmental stages for individual embryos, mapped onto the Atlas geometry (caudal view). Values represent superficial tissue changes, with values < 1 indicating compression, values = 1 isochoric deformation, and values > 1 expansion. Partial IFTs and the arterial pole are not present in the Atlas. Developmental stages represented by more than one embryo are included.

**Figure S6.**
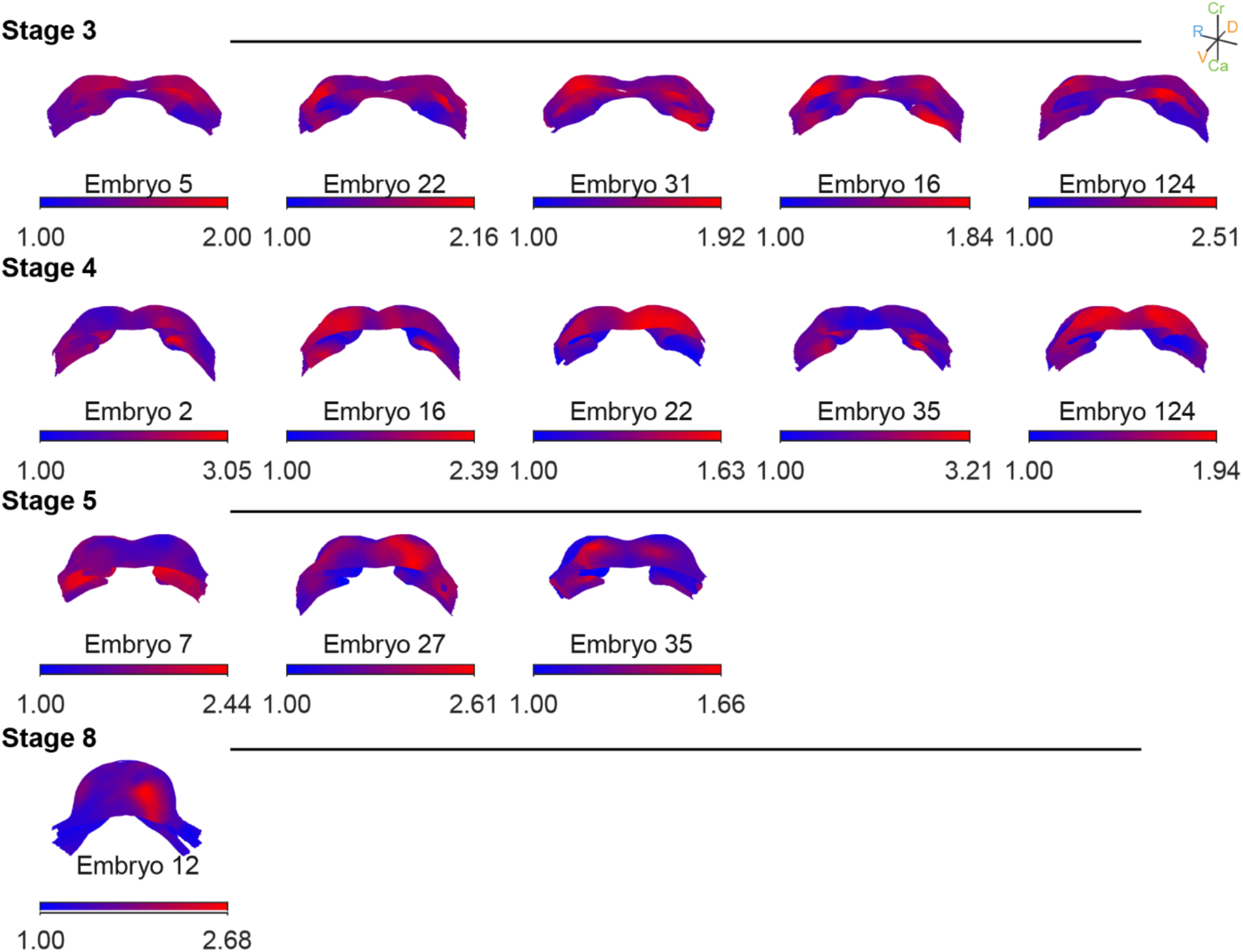
Mean anisotropy rate derived from stepwise deformations between consecutive developmental stages for individual embryos, mapped onto the Atlas geometry (caudal view). Values represent superficial tissue changes, with values = 1 isotropic deformation, and values > 1 anisotropic. Partial IFTs and the arterial pole are not present in the Atlas. Developmental stages represented by more than one embryo are included.

**Table S1.**
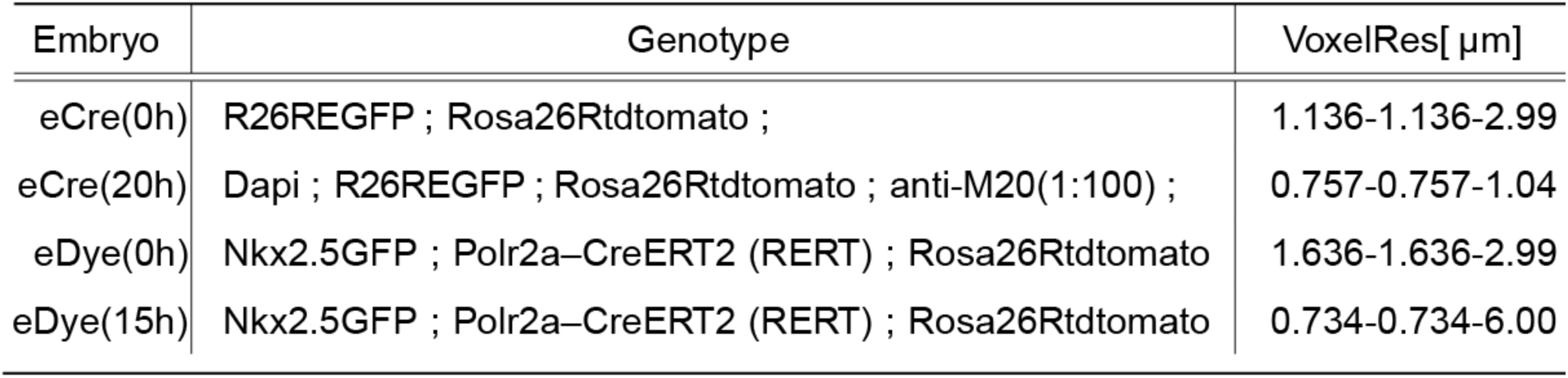
Dataset of TAT-Cre microinjection and cell Dye microinjection. eCre(20h) image was captured with Flash procedure (Messal et al.,2021)

**Table S2.** Dataset of cell dye microinjection experiments reporting labeling information for the right and left sides of the heart tube (HT). Embryos e1–e3 were used to measure D1, computed as a geodesic distance. Embryos e4–e5 were used to measure D2, computed as a Euclidean point-to-point distance between fixed landmarks. Both the landmarks and the ROI are available in the Mendeley dataset (Figure 5). Fold changes are reported between t(0) and t(end) over an approximately 10-hour time window.

| Embryo | Labeling |  | Fold change |
| --- | --- | --- | --- |
|  | Left | Right |  |
| e1_D1 | dil | diD | 0.23x |
| e2_D1 | dil | diD | 0.51x |
| e3_D1 | dil | diD | 0.77x |
| e4_D2 | dil | diD | 0.13x |
| e5_D2 | dil | diD | 0.29x |
| e6_D2 | diD | dil | 0.40x |

Video S1: In-silico Fate Map. The Dynamic Atlas is represented in its point-cloud version. At the cellular level, a yellow spot tracks one pseudo cell position throughout heart morphogenesis. At the tissue level, yellow spots illustrate the tissue deformation occurring during heart morphogenesis.

## Lead contact

Further information and requests for resources and code should be directed to and will be fulfilled by the lead contact, Miguel Torres and Domínguez Macías, Jorge Nicolás

## Materials availability

This study did not generate new unique reagents.

## Data and code availability

The authors declare that all the data supporting the findings of this study are available within the article and its supplemental files or from the corresponding author upon reasonable request. All data presented in this study, including raw and processed data have been deposited in a reserved Mendeley Data server at the following address:

- Mendeley Data: https://data.mendeley.com/preview/nd3kmj3cnx?a=a9cfaa55-3a74-4494-aeae-6cf0e90e5fa8

This paper does not report original code.

*In-silico* fate map is available in https://github.com/MorRaiola/BarrelHeartModel.git

